# Plasmids as donors of Insertion Sequence elements mediating colistin resistance in *Klebsiella pneumoniae*

**DOI:** 10.1101/2021.11.02.466793

**Authors:** Stephen Fordham, Anna Mantzouratou, Elizabeth Anne Sheridan

## Abstract

Colistin is a last resort antibiotic for the treatment of carbapenemase producing *Klebsiella pneumoniae* (CRKP). In line with rising colistin use worldwide, colistin resistant *Klebsiella pneumoniae* (*K. pneumoniae*) isolates have emerged. The disruption of *mgrB* by insertion sequences (ISs) has been widely reported worldwide representing a mechanism mediating colistin resistance. Evidence suggests plasmids encode mobilizable IS elements which preferentially integrate into the *mgrB* gene in *K. pneumoniae* causing gene inactivation and colistin resistance.

Recognised IS elements targeting *mgrB* include ISL3 (ISKpn25), IS5 (ISKpn26), ISKpn14 and IS903B-like elements. *K. pneumoniae* represents the single largest species carrying plasmids encoding these IS elements. For IS presence among species, 1000 BLASTn hits were downloaded and filtered for plasmids. Additionally, the top 120 BLASTn non-duplicate circularised plasmid contig hits for each IS element were typed for incompatibility (Inc) group and carbapenemase gene presence.

IS903B was found in 28 unique Inc groups, while ISKpn25 was largely carried by IncFIB(pQil) plasmids. ISKpn26 and ISKpn14 were most often found associated with IncFII(pHN7A8) plasmids. Of the 34 unique countries which contained any of the IS elements, ISKpn25 was identified from 26. ISKpn26, ISKpn14, and IS903B insertion sequences were identified from 89.3%, 44.9% and 23.9% plasmid samples from China.

Plasmids carrying ISKpn25, ISKpn14, and ISKpn26 IS elements are 12.18, 27.0, and 44.43 times more likely to carry carbapenemase genes relative to plasmids carrying the IS element IS903B. Moreover, plasmids carrying ISKpn26, ISKpn25, and ISKpn14 were 6.10, 28.82, and 31.47 times more likely to be sourced from a clinical environmental setting than the environment relative to IS903B IS harboring plasmid.

ISKpn25 present on IncFIB(pQil) sourced from clinical settings is established across multiple countries, while ISKpn26, ISKpn14, and IS903B appear most often in China. High carbapenemase presence in tandem with IS elements may help promote an extensively drug resistant profile in *K. pneumoniae* limiting already narrow therapeutic treatment options.

## 1. Introduction

Colistin serves as a last-resort antibiotic choice for the treatment of bacterial infections caused by carbapenemase producing *Klebsiella pneumoniae* (CPKP) and other gram-negative isolates. Rising colistin use has seen a corresponding increase in colistin resistance, especially during therapy and is an emerging global threat. From 2011 to 2015, in Italy, the rate of colistin resistance in *Klebsiella pneumoniae* (*K. pneumoniae*) increased from 36% to 50% (Giani *et al*., 2015). Separately, in Thailand, colistin resistance has been reported at 76.1% from a sample of 280 *K. pneumoniae* clinical isolates collected from 2014-17 (Eiamphungporn *et al*., 2018). Notably, colistin resistance due to disruption of the chromosomal *mgrB* gene in *K. pneumoniae* via the integration of insertion sequences (IS) has been widely reported from countries including: Lao PDR, Thailand, Nigeria, and France (Olaitan *et al*., 2014), Italy (Esposito *et al*., 2018), Greece (Giordano *et al*., 2018; Zhu *et al*., 2019; Hamel *et al*., 2020), Tunisia (Jaidane *et al*., 2018), Saudi Arabia (Zaman *et al*., 2018), Oman (Al-Farsi *et al*., 2019), Israel (Lalaoui *et al*., 2019), India (Shankar *et al*., 2019), Taiwan (Yang *et al*., 2020), USA (Macesic *et al*., 2020), and Malaysia (Yap *et al*., 2020).

Colistin binds to the lipopolysaccharide (LPS) component of the outer membrane of gram-negative bacteria. The cationic diaminobutyric acid (Dab) residues of colistin bind to anionic phosphate groups in the LPS. Colistin then destabilises both Mg^2+^ and Ca^2+^ divalent cations from the phosphate groups of LPS, disrupting the integrity of the membrane. Following membrane destabilisation, colistin binds to the lipid A moiety of LPS causing the derangement of the outer membrane (Falagas *et al*., 2005). Colistin resistance in *K. pneumoniae* is mediated by the modification of the LPS through the addition of 4-amino-4-deoxy-L-arabinose (l-Ara4N) to the phosphate groups of the lipid A moiety. l-Ara4N addition to LPS attenuates the affinity of colistin to LPS targets (Helander *et al*., 1996). The l-Ara4N induced modification of LPS is controlled by the products of the *pmrHFIJKLM* operon, positively regulated by the two component PhoQ/PhoP and PmrAB systems. MgrB, a product of the *mgrB* gene is a small transmembrane regulatory protein synthesized following the activation of the PhoQ/PhoP signalling cascade. MgrB interacts with the PhoQ sensor kinase exerting negative feedback on the PhoQ/PhoP system (Lippa *et al*., 2009). Insertional inactivation of *mgrB* prevents the down-regulation of the PhoQ/PhoP systems and represents a mechanism facilitating *de novo* acquired colistin resistance (Cannatelli *et al*., 2013).

Resistance to colistin frequently arises via the disruption of the *mgrB* gene by insertion sequences (ISs) in *K. pneumoniae*. IS-mediated *mgrB* gene disruption can represent a significant cause of colistin induced resistance in *K. pneumoniae*. Across two independent samples of 31 and 49 clinically isolated colistin resistant *K. pneumoniae* from Taiwan, 34.7% (*n*=17/49), and 32.2% (*n*=10/31) carried IS elements in the *mgrB* gene (Yang *et al*., 2020; Berglund *et al*., 2018), while 30% (*n*=6/20) clinical colistin resistant *K. pneumoniae* isolates from Iran carried IS elements insertion, either IS5-like or IS1-like in *mgrB* (Haeili *et al*., 2017). More strikingly, from a sample of 11 clinical colistin resistant *K. pneumoniae* isolates from China, 9 carried IS elements in the *mgrB gene* (Yan *et al*., 2021), while 93.75% (*n*=15/16) clinical isolates from Greece harboured ISKpn26-like element disruption in *mgrB* (Zhu *et al*., 2019). ISs elements including ISKpn25, ISKpn26, IS903B and ISKpn14 are common IS elements targeting genes involved in colistin resistance (Haeili *et al*., 2017; Berglund *et al*., 2018; Di Tella *et al*., 2019; Yang *et al*., 2020; Yan *et al*., 2021). Insertional inactivation of *mgrB* represents a prominent mechanism mediating the emergence of colistin resistance in *K. pnuemoniae*.

*De novo* colistin resistance may occur through the transposition of ISs from plasmids into chromosomal colistin associated genes. Across a sample of 29 and 19 colistin resistant *K. pneumoniae* isolates from Italy and Greece, 2 clonally related clusters; 2 Italian ST512 isolates and 8 Greek ST258 *K. pneumoniae* isolates harboured the complete copy of ISKpn25 inserted at nucleotide position 133 of the *mgrB* gene (Giordano *et al*., 2018). The same ISKpn25 was located on pKpQIL-like plasmids from these samples, indeed the ISKpn25 on pKpQIL plasmids and the ISKpn25 inserted into the *mgrB* gene share a 100% match between the 8154 nucleotides. Furthermore, in 2 ST258 and 2 ST512 *K. pneumoniae* isolates from Greece and Italy, the IS5-element derived from pKpQIL-like plasmids, was found inserted into nucleotide position 75 of the *mgrB* gene (Giordano *et al*., 2018). Nucleotide position 75 represents a hotspot for IS5 element insertion among clonally unrelated *K. pneumoniae* isolates (Cannatelli *et al*., 2013; Poirel *et al*., 2015). Mobilisation of IS5 elements from plasmids into the *mgrB* gene of colistin resistant *K. pneumoniae* has been speculated owing to the similarly between the IS5 element inserted into *mgrB* and the IS5 element present on *K. pneumoniae* carbapenemase (KPC)-encoding and other Gram-negative bacterial plasmids (Cannatelli *et al*., 2013; Hala *et al*., 2019; Azam *et al*., 2021), and endogenous presence of IS5-like elements in the genome of colistin resistant *K. pneumoniae* (Choi *et al*., 2020).

Further evidence for the involvement of plasmids as donors for IS elements is provided by a recent cloning assay. IS elements including ISKpn26, ISKpn14, and IS903B cloned into a plasmid vector and transformed into a colistin susceptible *K. pnuemoniae* isolate increased the frequency of colistin resistance *K. pneumoniae* isolates. Notably, for the plasmid vector carrying IS903B, colistin induced stress was responsible for IS mobilization (Yang *et al*., 2020). Furthermore, a *Caenorhabditis elegans* killing assay model revealed nematodes fed with *K. pneumoniae* isolates harbouring plasmids carrying ISKpn26 were associated with a significantly reduced lifespan and higher death risk during colistin treatment relative to nematodes fed with non-IS plasmid carrying *E*.*coli* and *K. pneumoniae* isolates, thus confirming the role of plasmids as donors for IS elements mediating colistin resistance (Yang *et al*., 2020).

Beyond gene insertion, IS elements can integrate into the promoter regions of chromosomal colistin-resistance associated genes. The IS1R IS element sourced from a plasmid has been shown to integrate into the promoter region of *mgrB* mediating the emergence of a *de novo* colistin resistant phenotype (Antonelli *et al*., 2017), indeed IS1-like element insertion has been independently reported repeatedly in the promoter region of *mgrB* (Haeili *et al*., 2017; Berglund *et al*., 2018). For IS1382-like and IS1-like elements disrupting *mgrB* in *K. pneumoniae*, BLASTn searches revealed their presence only in respective isolates with disruptions in *mgrB*, notably absent from any other chromosomal location, thereby indicating a likely plasmid source as opposed to transposition from a chromosomal source (Jaidane *et al*., 2018).

Colistin resistance may arise through the transposition of ISs from source IS containing plasmids preferentially targeting specific *mgrB* regions for recombination. The dissemination of IS elements that transpose into the same position of the *mgrB* gene may represent a mechanism mediating the observed colistin resistance in both clonally related and unrelated *K. pneumoniae* isolates. IS elements are frequently reported in *K. pneumoniae* colistin-associated chromosomal resistant genes. In order to ascertain IS element prevalence in *K. pneumoniae*, IS elements which have been shown to disrupt the *mgrB* gene; namely ISKpn25, ISKpn26, ISKpn14 and IS903B were investigated. Species IS prevalence among 1000 BLASTn hits was assessed. Separately, to support pathogen surveillance, plasmid incompatibility typing (Inc) for the top 120 BLASTn plasmid hits were performed to determine the respective plasmid incompatibility group associated with each IS element. In addition, metadata for the same plasmid samples was gathered to determine dual carbapenemase gene prevalence, plasmid size, and species, country, and isolation source in order to elucidate the emerging clinical threat posed by plasmid bearing IS elements.

## 2. Method

4 insertion sequence reference elements were derived from the online ISfinder tool (https://www-is.biotoul.fr). The IS elements include: IS1 (ISKpn14), ISL3 (ISKpn25), IS5 (ISKpn26), and IS903B insertion sequences (Table 1a). Each IS element was blasted using the NCBI nucleotide blast online tool, BLASTn. To assess IS proportional assignment between bacterial species, the max target sequences parameter was increased from the default of 100 to 1000 for each IS element. The description table for each BLASTn hit was downloaded and filtered for plasmids to ascertain plasmid samples encoding IS elements. For ISKpn25 which returned many low identity and coverage hits, plasmid hits above a minimum threshold; percentage identity ≥95% was used. This threshold was chosen to differentiate ISKpn25 from ISKpn26 which shares 91% nucleotide identity (https://www-is.biotoul.fr). The relative proportion of IS elements among *K. pneumoniae* samples against other bacterial species was calculated and plotted.

**Table 1a.** Reference IS elements soured from IS finder with their associated length, open reading frame, open reading frame function and accession numbers. **\*1= Transposase, 2= Putative restriction endonuclease S subunit, 3= Hypothetical protein, 4 = Putative type I restriction-modification system DNA methylase**

| *Reference IS element* | Length (bp) | Accession Number | Open Reading Frame(s) | Open Reading Frame function |
| --- | --- | --- | --- | --- |
| *IS1 (ISKpn14)* | 768 | CP000649 | 91 (56-331)<br>174 (229-753)<br>274 (56-753) | *1,<br>1,<br>1 |
| *ISL3 (ISKpn25)* | 8154 | NC_009650 | 684 (107-2161)<br>438 (2158-3474)<br>1085 (3475-6732)<br>422 (6737-8005) | *1,<br>2,<br>3,<br>4 |
| *IS5 (ISKpn26)* | 1196 | NC_016845 | 326 (69-1049) | *1 |
| *IS903B* | 1057 | X02527 | 307 (78-1001)<br>70 (672-460) | *3,<br>1 |

For each IS element, the top 120 non-duplicate circularized plasmid hits harbouring IS elements were downloaded, and the Fasta contigs incompatibility (Inc) typed using PlasmidFinder (updated September, 2020). Each plasmid contig carrying IS elements was also scanned for carbapenemase genes with a predicted *in silico* resistance phenotype to the carbapenem, meropenem. Metadata pertaining to each sample, including accession number, species and isolation source, country of origin, and plasmid size was recorded by accessing available Biosample information from the BLASTn hits from NCBI. Results were tabulated and analysed using Python and are available in Supplementary Table s1. Figures were produced using both the Matplotlib and Seaborn libraries in Python. For statistical analyses between the IS elements and the presence of carbapenemase genes/sample source, the chi-square (χ^2^) test of independence was used with an α value of 0.05 as the significance level. To correct against multiple comparisons between each IS element, *p*-values were corrected using the False Discovery rate (FDR) method. To measure the strength of association between carbapenemase gene prevalence/clinical source isolation against different IS elements, both Cramer’s V and the odds ratio (OR) was calculated for each pair of IS elements. Statistical analyses were conducted using IBM SPSS Statistics v28.0.0. The criterion outlined in Table 1b was used for Cramér’s V effect size characterisation (IBM, 2021). The method workflow is shown in Figure 1.

**Table 1b.** Cramer’s V is an effect size measurement used for the chi-square test of independence. Cramer’s V measure the strength of association between two categorical fields.

| *Effect size (ES)* | Interpretation |
| --- | --- |
| ES ≤ 0.2 | Fields are weakly associated |
| 0.2 < ES ≤ 0.6 | Fields are moderated associated |
| ES > 0.6 | Fields are strongly associated |

**Figure 1.**
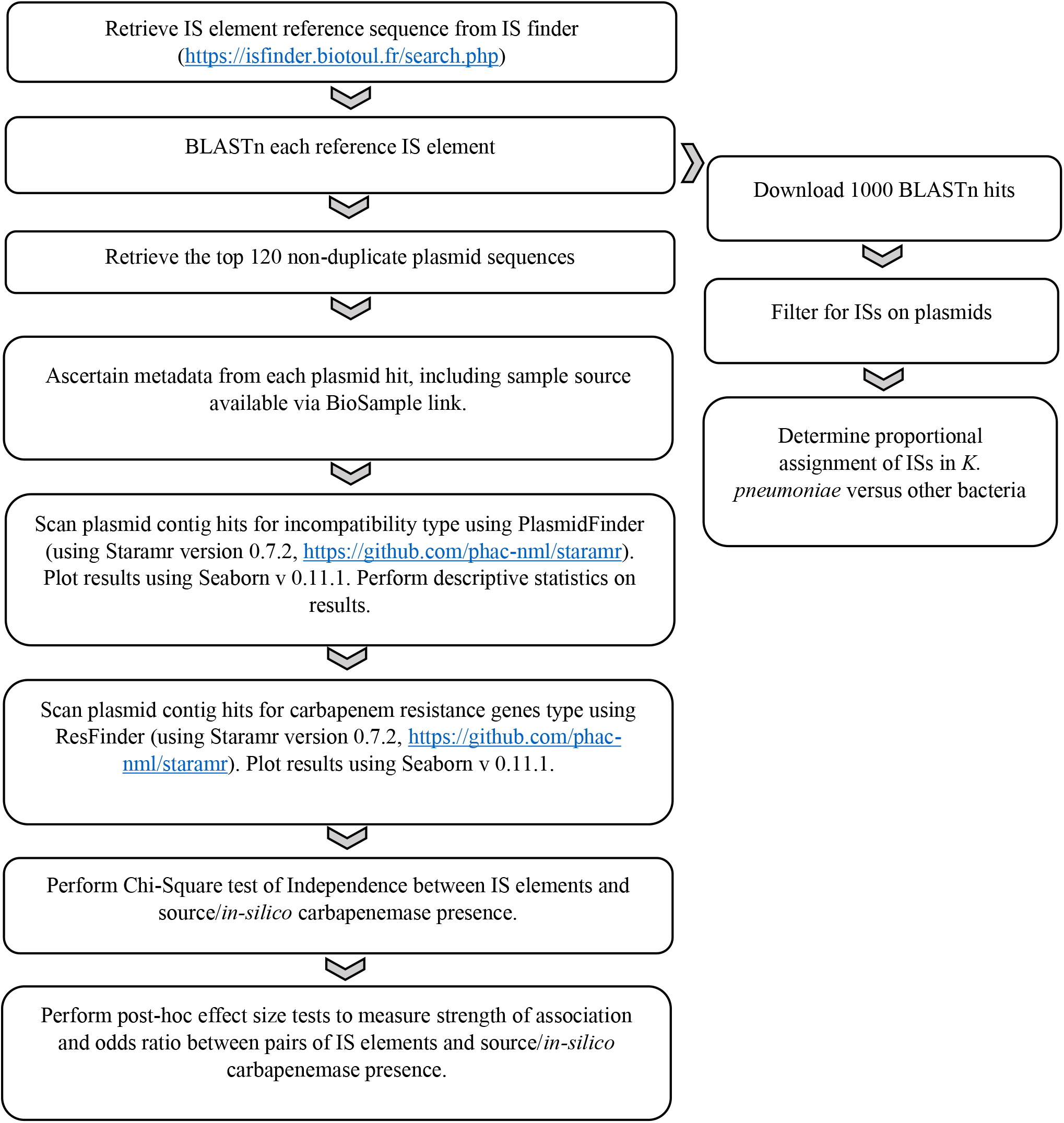
Method Workflow.

For clinical sample classification, Biosample data labelled as: blood, urine, human hospital, rectum, feces, sputum, necrotic tissue, throat, wound swab, clinical sample, hospital environment, ascites, bile, tissue sample, wound, abdominal pus, secreta, stool, lymphocele, intestine fluid, groin, abscess, bronchoalveolar lavage and ulcers were classified as clinically derived samples, while samples labelled as environment, pig, horse, intestines (animal), sample milk, sewage, *Manis javanica*, waste treatment plant, food, equine body fluid, and dog were classified as environmentally derived samples in our analysis.

Biosamples termed wastewater, wastewater influent sample and wastewater effluent samples are not categorized as either clinical samples or environmental samples. A total of 21 samples fit into this third undetermined category. These samples do not feature in either the frequency data pool used in both the Chi-Square test of Independence between IS elements and sample source, and subsequent post-hoc Cramer’s V and odds ratio association effect size tests. Sample source for each plasmid can be found in Supplementary Table s1.

## 3. Results

### 3.1 Insertion Sequence element pervasiveness in *K. pneumoniae*

*K. pneumoniae* represents the dominant species harbouring IS elements; ISKpn25, ISKpn26, ISKpn14 and IS903B on plasmids. Across the sample of plasmids encoding ISKpn25 (*n*=173, mean query coverage: 99.85%, standard deviation: 0.92%, mean identity: 99.87%, standard deviation: 0.73% against reference NC_009650), *K. pneumoniae* constituted 90.17% (*n*=156) of all samples carrying ISKpn25 (Figure 2a, ISKpn14). In addition, *K. pneumoniae* carried the IS element ISKpn25 at a 31.2-fold higher rate than the second most abundant bacterium encoding ISKpn25 on plasmids, namely *E. coli* (Figure 2b, ISKpn25).

**Figure 2.**
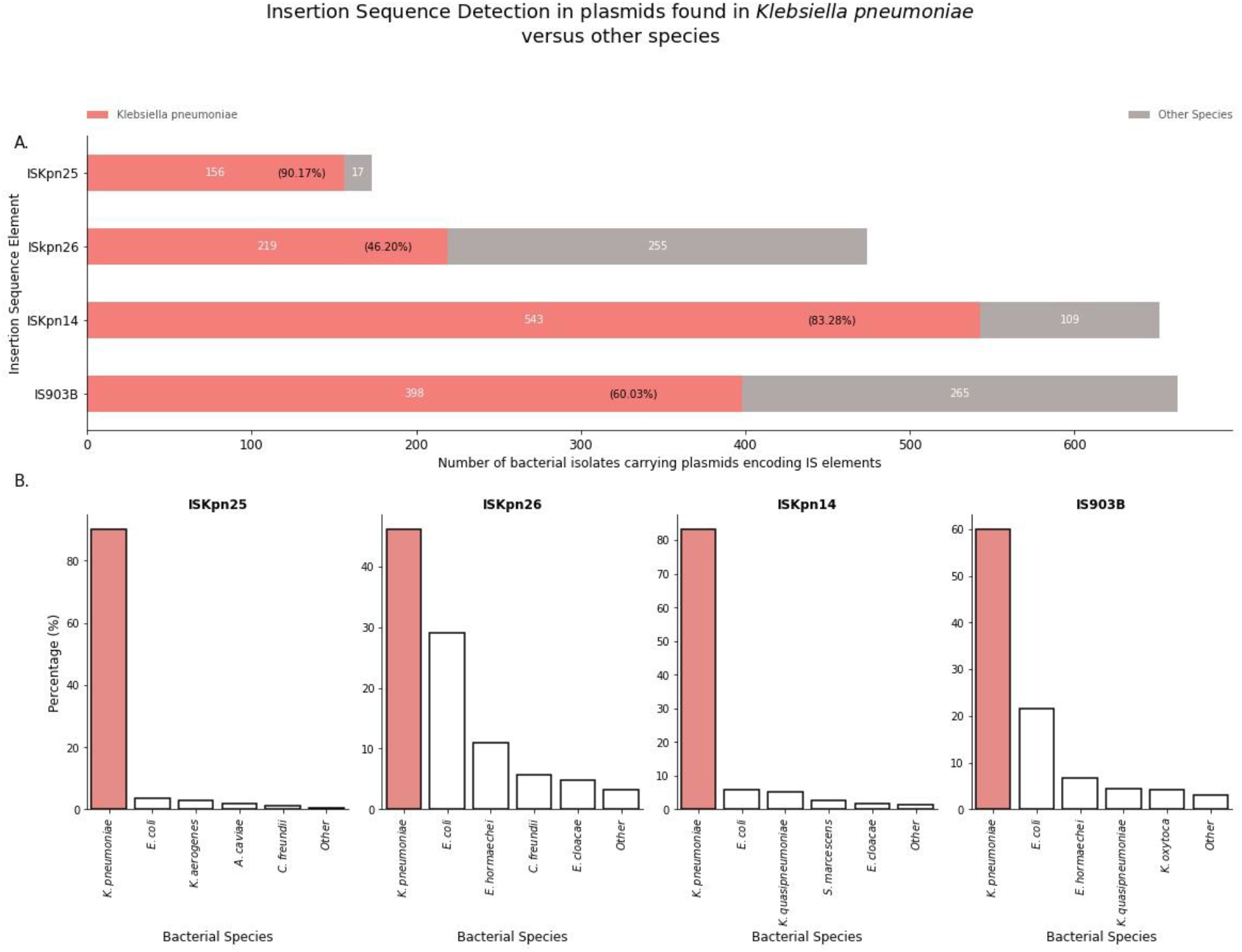
Proportional assignment of IS elements ISKpn25, ISKpn26, ISKpn14 and IS903B among *Klebsiella pneumoniae* and other bacterial species. A. The max target sequences parameter on the NCBI BLASTn tool (https://blast.ncbi.nlm.nih.gov) was adjusted from the default of 100 to 1000, to gain insight into IS proportional assignment among species. For each IS element, *K. pneumoniae* plasmid hits are shown as red horizontal bars with bacterial counts indicated in white, percentages in parentheses in black. B. *K. pneumoniae* bars shaded in red depict species percentage; the closest 4 species are given a separate white bar, while the reminding species are grouped into a single white bar labelled other.

*K. pneumoniae* also represents the most common species carrying the IS element ISKpn26 on plasmids. Across the sample of plasmids encoding ISKpn26 (*n*=474, mean query coverage: 100%, mean identity: 99.78%, standard deviation: 0.13% against reference NC_016845), *K. pneumoniae* comprised 46.20% of plasmid samples carrying ISKpn26, a rate 4.2-fold higher than *E*.*coli* (Figure 2ab, ISKpn26).

Across a sample of 652 plasmid samples encoding ISKpn14 (mean query coverage: 100%, mean identity: 99.98%, standard deviation: 0.11% against reference CP000649), *K. pneumoniae* was the principal bacterium carrying ISKpn14 on plasmid samples, responsible for 82.3% (*n*=543) of ISKpn14 plasmid samples (Figure 2a, ISKpn14). In fact, *K. pneumoniae* carried ISKpn14 at a rate 15.9-fold higher relative to *E. coli*, the second most abundant ISKpn14-carrying species (Figure 2b, ISKpn26).

Finally, in line with observations for the other three IS elements, *K. pneumoniae* was the dominant species encoding IS903B-like IS elements on plasmids. IS903B-like ISs were found on 60.03% (*n*=398) of IS903B-plasmid containing samples (mean query coverage: 99.99%, standard deviation: 0.03%, mean identity: 98.72%, standard deviation: 0.21% against reference X02527). *K. pneumoniae* harboured IS903B at an 8.84-fold higher rate relative to *E*.*coli*, the second most abundant species encoding IS903B (Figure 2b, IS903B).

### 3.2. Insertion Sequence element stratification among plasmid incompatibility groups

The four IS elements follow a different stratification pattern among plasmid incompatibility groups (Inc). The IS element, IS903B is the most diverse plasmid host range element, present in 28 unique Inc groups or Inc groups representing fusion plasmids. Relative to ISKpn14, ISKpn25, and ISKpn26, IS903B was present in 1.42, 1.83, and 2.33-fold higher unique Inc groups, respectively (Figure 3). Despite the broad host range of IS903B among various unique plasmid Inc groups, 16/28 represented a single instance of the IS element associated with a unique plasmid Inc group. Indeed, for the other IS elements investigated, a single Inc group was found in 9, 7 and 4 instances for the IS elements ISKpn14, ISKpn26, and ISKpn25, respectively. To determine the dominant Inc groups associated with the IS elements, the dataset was filtered for ≥ 5 occurrences of the same Inc group. Here, 3, 3, 4, and 6 unique Inc groups were identified in plasmids harbouring ISKpn25, ISKpn26, IS903B and ISKpn14 elements, respectively. Notably, the IncFIB(pQil) Inc group was present in 88/120 samples harbouring ISKpn25. Furthermore, the same IncFIB(pQil) Inc group in addition to either IncR or IncFII(K) was found in an additional 5 and 7 plasmids encoding the insertion element ISKpn25. From the top 120 BLASTn hits, the ISKpn25 element is disproportionately identified in plasmid belonging to the IncFIB(pQil) replicon family. Likewise, plasmids harbouring the IS elements, ISKpn26 and ISKpn14 follow a pronounced partitioning among Inc groups. The ISKpn26 elements were associated with IncFII(pHN7A8) encoding plasmids. Here, IncFII(pHN7A8) was found in *n*=36/120 plasmids while *n*=44/120 plasmids carried both IncFII(pHN7A8) and the IncR group. ISKpn14 was associated with the IncHI2A and IncHI2 fusion group. In contrast, the IS element IS903B is associated with various plasmid Inc groups including IncHI2A + IncHI2, IncFIB(K), IncR and RepA, as shown in Figure 3. Taken together, IS elements with demonstrable ability to disrupt chromosomal colistin-associated genes are present in a large number, namely 61 unique Inc groups from a sample size of 480 BLASTed IS elements. Despite this, specific Inc groups are disproportionately associated with particular IS elements. Typing plasmids may help to yield the relationship between IS elements and their partner plasmid families.

**Figure 3.**
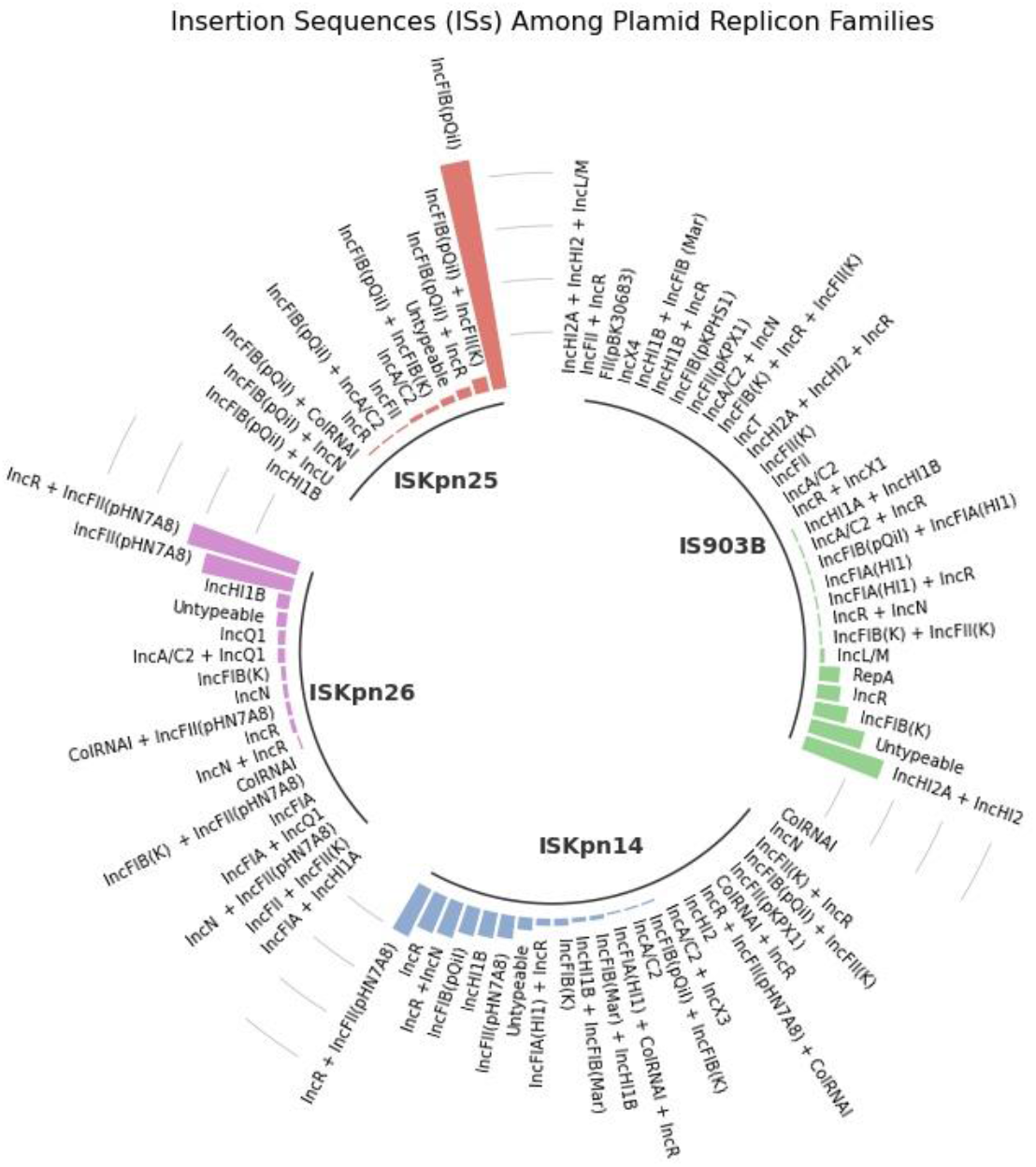
Insertion Sequences (ISs) stratification among plasmid replicon families. IS elements identified in plasmids belonging to various plasmid Incompatibility (Inc) groups. The circular bar plot schematic represents the results of the top 120 IS BLASTn hits for each insertion element present in plasmids (Accession numbers for IS elements listed in method, Table 1a). Relative to ISKpn25 colored red, identified in 12 unique Inc groups, ISKpn26 colored purple, is present in 17 unique Inc groups, 1.42-fold higher), ISKpn14 colored blue, is present in 22 unique Inc groups, 1.83-fold higher, while IS903B, colored green, is the most diverse plasmid host-range IS element, present in 28 unique Inc groups, with a 2.33-fold higher plasmid replicon count. The dominant Inc groups per insertion sequence are IncFIB(pQil) for ISKpn25 IS elements, the fusion plasmid IncHI2A + IncHI2 for IS903B IS elements and the fusion plasmid IncR + IncFII(pHN7A8) for ISKpn26 and ISKpn14 IS elements, respectively. Plasmids which were classified as untypeable were not regarded as a unique Inc group.

The four IS elements are associated with diverse plasmids with a varied size ranging from 10,159bp through to 490,750bp (Figure 4). Notably, of the 34 unique countries which contained any of the IS elements, ISKpn25 was identified from 26. India, USA, and Italy were the country of origin for 16.6%, 13.3%, and 9.2% of ISKpn25 insertion element plasmid samples. In contrast, typed ISKpn26 elements are associated with IncFII(pHN7A8) Inc groups, and are largely restricted to China, 89.3% (n=107/120), with a further 7 samples identified from Taiwan. Similar to ISKpn26 encoding elements, the largest proportion of typed samples encoding ISKpn14, 44.9%, were isolated from China. Finally, China represents the single largest country harbouring IS903B samples, with a 23.9% proportional assignment, however IS903B is also found at similar levels in the USA (18.8%), Australia (18.8%), and the UK (14.5%), suggesting IS903B is established internationally. Across the four IS elements, 74.16% (*n*=356/480) typed plasmid samples were sourced from clinical samples. China is the largest country harbouring ISKpn26 (*n*=73/83, 87.9%) and ISKpn14 (*n*=53/107, 49.5%) clinical samples, while China is the second largest contributor for IS903B derived clinical samples, (*n*=18/68, 26.5%). In contrast, the clinically sourced ISKpn25 samples are derived from 24 countries, although India contributes 20.6% of these samples. Taken together, IS elements present in plasmids are endemic in China, established in other countries and both the environment and clinical settings may represent sources for IS elements. Notably, the environment may serve as a potential origin for plasmid encoded IS903B samples whereby 19.05% (*n*=20/105) of plasmids encoding this IS element are derived from environmental sources, and a further 16.2% (*n*=17/105) did not disclose an isolation source. Collectively, these relationships are summarised in Figure 4c.

**Figure 4.**
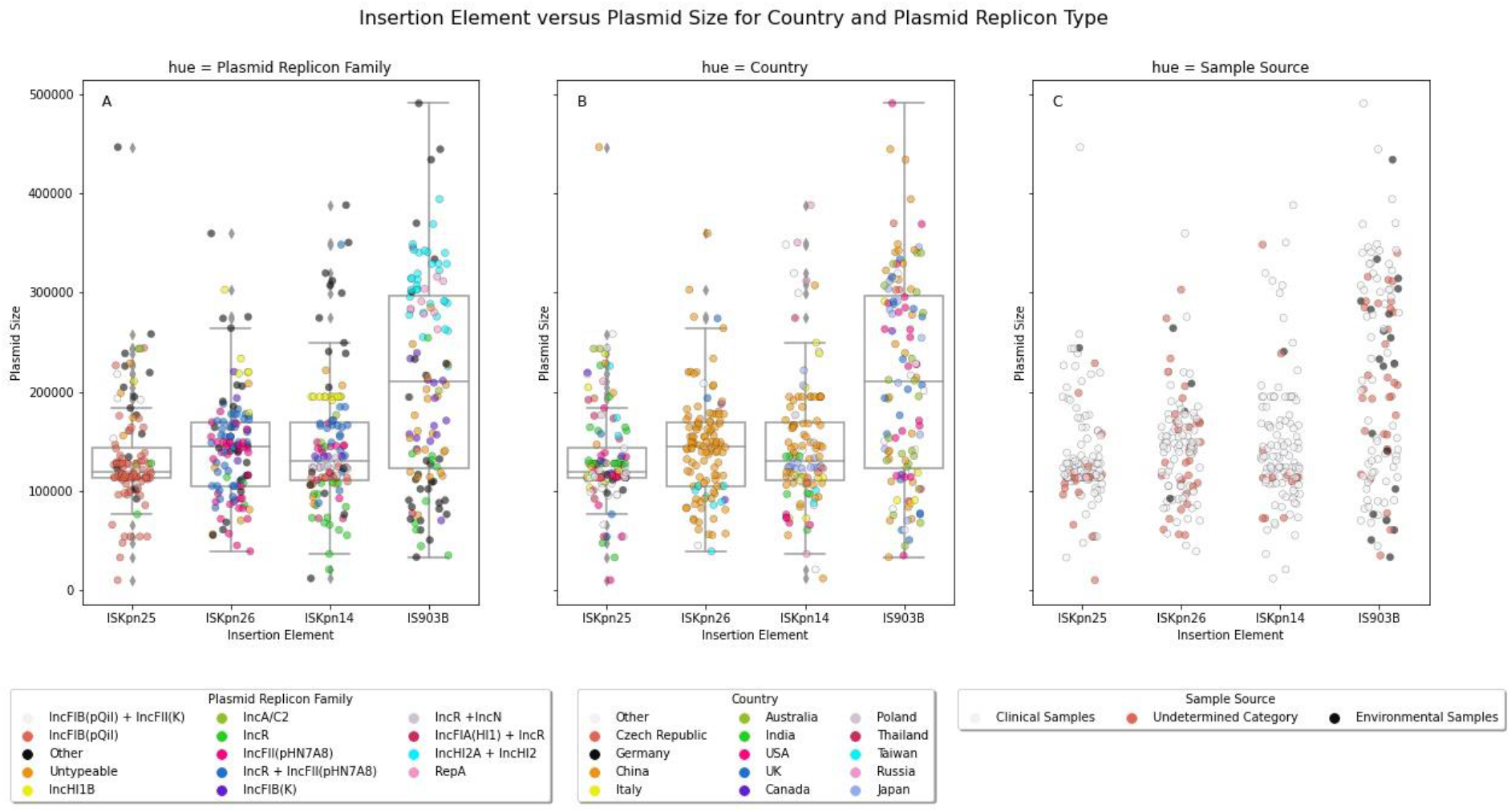
Insertion element stratification for ISKpn25, ISKpn26, ISKpn14 and IS903B. A. Plasmid replicon family versus plasmid size for the four IS elements. B. Country stratification versus plasmid size. C. Source sample stratification. Each IS element included 120 non-duplicate plasmid samples. Plot produced using the Seaborn library in Python.

### 3.3. Plasmids encoding both IS elements and carbapenemase genes

Plasmids carrying IS elements were additionally investigated for the presence of carbapenemase genes. A marked stratification pattern exists for the various IS elements and carbapenemase genes. For plasmids harbouring either ISKpn26, ISKpn14 or ISKpn25, 82.5%, 75% and 57.5% also carried carbapenemase genes with a predicted *in silico* resistance phenotype against the carbapenem, meropenem. In contrast, only 10% of plasmids carrying the IS element IS903B also carried carbapenemase genes (Figure 5). The chi-square (χ^2^) test of independence revealed a significant difference between carbapenemase gene distribution between the 4 IS elements, χ^2^(3, *N* = 480) = 155.12, p = <.001. Post-hoc analysis revealed a significant difference between IS903B, and the three other IS elements. Relative to IS903B, carbapenemase gene distribution was significantly different relative to ISKpn25, ISKpn26, and ISKpn14: IS903B versus ISKpn25 (*X*^2^, p<0.001), IS903B versus ISKpn14 (*X*^2^, p<0.001), and IS903B versus ISKpn26 (*X*^2^, p<0.001) Results are summarised in Table 2a-c. A statistically significant difference between the distribution of sample source for IS elements was also observed, χ^2^(3, *N* = 382) = 46.97, p = <.001. A significant difference between the distribution of sample source for IS903B against ISKpn14 (*X*^2^, p<0.001), ISKpn26 (*X*^2^, p<0.001) and ISKpn25 (*X*^2^, p<0.001) was also observed. Results are summarised in Table 2d.

**Figure 5.**
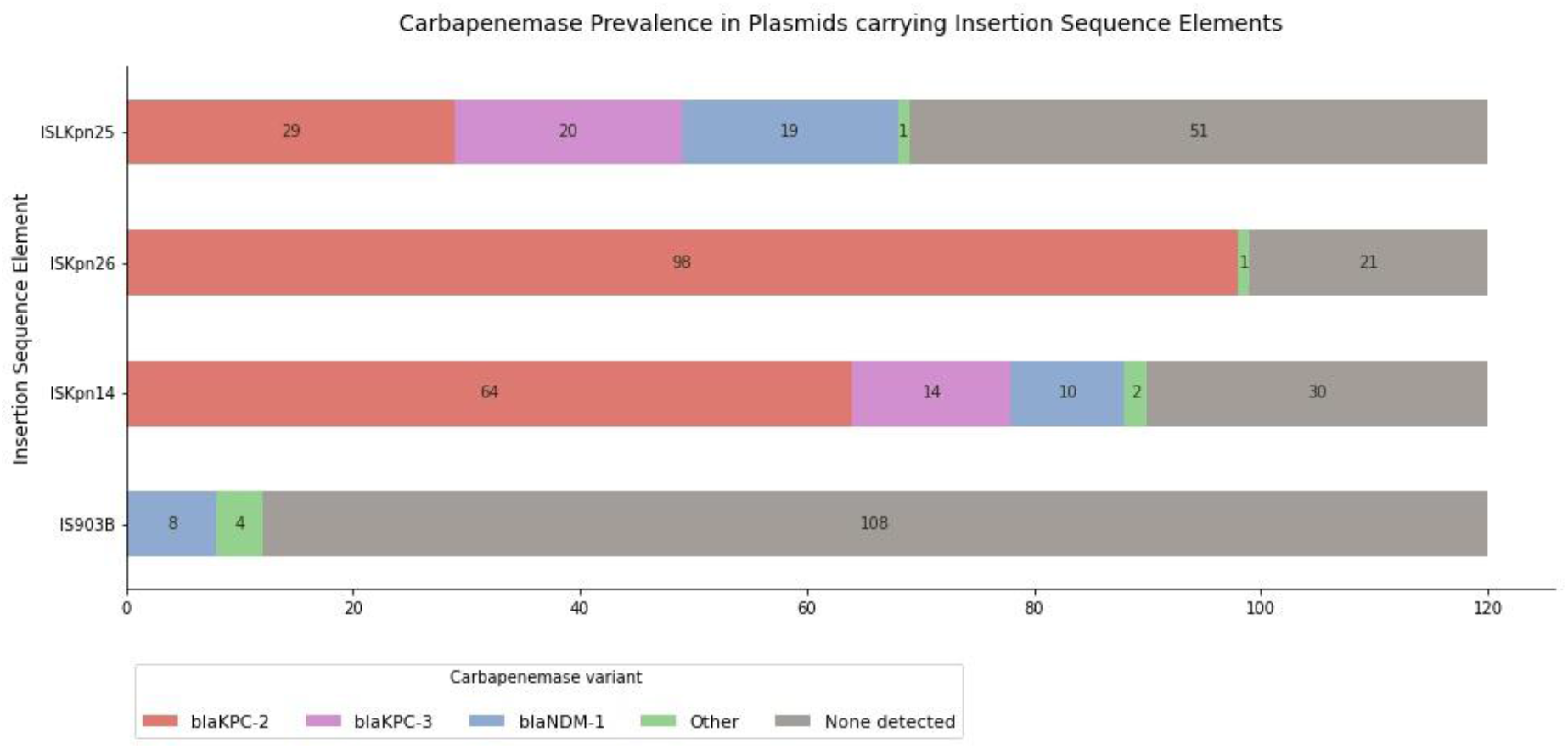
Carbapenemase prevalence in plasmids carrying IS elements. Plasmids encoding IS elements were also investigated for the presence of carbapenemase genes by scanning plasmid contigs against the ResFinder 4.0.0 database. The listed carbapenemase variants *bla*_KPC-2_, *bla*_KPC-3_, *bla*_NDM-1_, had respective counts ≥1, and are shown as individual categories. The other category comprises carbapenemase genes variants which were grouped together based on their singular appearance with IS elements. These other carbapenemase genes include: *bla*_GES-5_, *bla*_IMP26_, *bla*_KPC-6_, *bla*_NDM-4_, *bla*_NDM-5_, *bla*_NDM-7_, *bla*_IMP-4_, *bla*_IMP26_, and _*bla*VIM-5_. Carbapenemase gene detection was based on the predicted *in silico* resistance phenotype to the carbapenem, meropenem.

**Table 2a.** The association between *in silico* carbapenemase gene presence and IS elements: IS903 versus ISKpn25, ISKpn14, and IS5 (ISKpn26), significant results (p<0.05) presented only.

| *IS Element* | **IS903B** |  |  |  |
| --- | --- | --- | --- | --- |
| | Cramer's V ($\phi_c$) | Odds ratio | *P* value | Pearson's $\chi^2$ |
| *ISKpn25* | 0.502 | 12.176 | <.001 | $X^2 (1, N = 240) = 60.545$ |
| *ISKpn14* | 0.657 | 27.000 | <.001 | $X^2 (1, N = 240) = 103.734$ |
| *IS5 (ISKpn26)* | 0.727 | 44.429 | <0.01 | $X^2 (1, N = 249) = 126.864$ |

**Table 2b.**
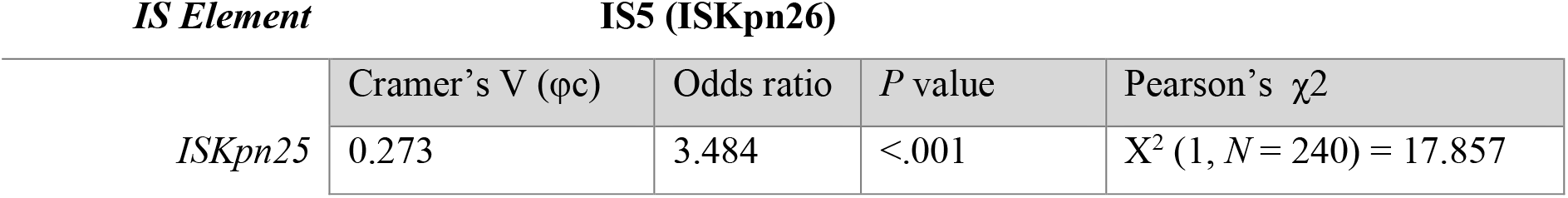
The association between *in silico* carbapenemase gene presence and IS element: IS5 (ISKpn26) versus ISKpn25.

| *IS Element* | **IS5 (ISKpn26)** |  |  |  |
| --- | --- | --- | --- | --- |
| | Cramer's V ($\phi_c$) | Odds ratio | *P* value | Pearson's $\chi^2$ |
| *ISKpn25* | 0.273 | 3.484 | <.001 | $X^2 (1, N = 240) = 17.857$ |

**Table 2c.** The association between *in silico* carbapenemase gene presence and IS element: ISKpn14 versus ISKpn25.

| *IS Element* | **ISKpn14** |  |  |  |
| --- | --- | --- | --- | --- |
| | Cramer's V ($\phi_c$) | Odds ratio | *P* value | Pearson's $\chi^2$ |
| *ISKpn25* | 0.165 | 2.043 | <.001 | $X^2 (1, N = 236) = 6.461$ |

**Table 2d.** The association between clinical source and IS elements. IS903 versus ISKpn25, ISKpn14, and IS5 (ISKpn26), significant results (p<0.05) presented only.

| *IS Element* | **IS903B** |  |  |  |
| --- | --- | --- | --- | --- |
| | Cramer's V ($\phi_c$) | Odds ratio | *P* value | Pearson's $\chi^2$ |
| *ISKpn25* | 0.343 | 28.824 | <.001 | $X^2 (1, N = 187) = 22.041$ |
| *ISKpn14* | 0.351 | 31.471 | <.001 | $X^2 (1, N = 196) = 24.992$ |
| *IS5 (ISKpn26)* | 0.264 | 6.103 | <0.01 | $X^2 (1, N = 175) = 12.151$ |

Cramer’s V (φc) reveals a strong or moderate association, (0.2 < ES ≤ 0.6/ ES > 0.6) between IS elements (IS903b-paired IS elements) and *in silico* carbapenemase gene presence (Table 1a). The odds ratio reveals plasmids carrying ISKpn25, ISKpn14, and ISKpn26 IS elements are 12.18, 27.0, and 44.43 times more likely to carry carbapenemase genes relative to plasmids carrying the IS element IS903B. Results are summarized in Table 2a. In addition, Cramer’s V determined a moderate effect size (φc = 0.271; 0.2 < ES ≤ 0.6) between the paired IS elements ISKpn26, ISKpn25 and *in-silico* carbapenemase gene presence, where ISKpn26 is 3.48 times more likely to be positive for carbapenemase genes relative to ISKpn25 (Table 2b). A weak effect size was recorded between the paired IS elements ISKpn14, ISKpn25 and *in-silico* carbapenemase gene presence; φc = 0.165; ES ≤ 0.2, where ISKpn14 is 2.2 times more likely to carry carbapenemase genes (Table 2c).

Cramer’s V also revealed a moderate association between IS903b-paired IS elements and source isolation. ISKpn26, ISKpn25, and ISKpn14 were 6.103, 28.824, and 31.471 times more likely to be sourced from a clinical environment than the environment relative to IS903B IS harboring plasmids. These relationships are summarized in Table 2d.

## 4. Discussion

*K. pneumoniae* represents a key reservoir species harbouring plasmids encoding IS elements which can disrupt the *mgrB* gene leading to colistin resistance. The IS elements, ISKpn25, ISKpn26, ISKpn14 have a majority percentage prevalence within *K. pneumoniae* across the sampled dataset, while *K. pneumoniae* represents the single largest species group carrying the IS903B element. The IncFIB(pQil) plasmid Inc group was associated with the IS element ISKpn25. This association has been previously observed in carbapenem and colistin resistant *K. pneumoniae* isolates derived from independent samples from both Greece and Italy (Giordano *et al*., 2018). The ISKpn25 containing IncFIB(pQil) plasmid appears highly successful, present in up to 26 countries from the sampled data, while 7 countries harboring ≥ 5 IncFIB(pQil) ISKpn25 samples have been detected. These countries include India, USA, Italy, Thailand, Germany, Canada, and Poland. From the countries with ≥ 5 samples, 94.2% (*n*=64/69), IncFIB(pQil) ISKpn25 samples are derived from clinical sources, suggesting the IS element ISKpn25 associated with IncFIB(pQil) plasmids are well established in clinical settings from geographically distinct areas. This may present a serious clinical threat, as selective colistin pressure in the hospital environment has been linked to the genetic transposition of IS elements in *K. pneumoniae* (Berglund *et al*., 2018; Yang *et al*., 2020; Yan et al., 2021). In this scenario, there may be few barriers to prevent inducible colistin resistance among bacterial isolates encoding ISKpn25 on their plasmids.

In contrast, IS elements ISKpn14 and ISKpn26 are most often identified from clinical samples from China. These results agree with recent molecular analyses which reveal the presence of ISKpn14 as the principle IS element disrupting the *mgrB* gene of colistin resistant clinical *K. pneumoniae* isolates from 6 hospitals across China (Yan *et al*., 2021). ISKpn26 is associated with IncFII or IncFII and IncR fusion plasmids. Interestingly, IncFII and IncR fusion plasmids, and IncR plasmids are the most commonly identified plasmid Inc groups from ISKpn14 carrying plasmids, suggesting these plasmid Inc groups are receptive towards IS element uptake and maintenance. In addition, both ISKpn26 and ISKpn14 containing plasmids also carried a high percentage of carbapenem resistant genes, a feature also detected in ISKpn14/ ISKpn26 *mgrB* disrupted *K. pneumoniae* isolates from China (Yan *et al*., 2021). This may represent a worrisome clinical threat, based on evidence that CPKP isolates are successful in clinical settings. *K. pneumoniae* isolates are more likely to have a genetically nearest neighbor (GNN) from the same hospital if they harbor carbapenemase genes (David *et al*., 2019). The clonal success imposed by carbapenemase presence may help disseminate IS elements. The combination of carbapenemase genes in tandem with IS may produce *K. pneumoniae* isolates which are difficult to treat. Furthermore, The IncFII typed pHN7A8 plasmid was present in 70.8% (*n*=85/120) and 26.7% (*n*=32/120) plasmids carrying ISKpn26 and ISKpn14, respectively.

Conjugation assays reveal IncFII -pHN7A8-like plasmids are highly transferable (Sennati *et al*., 2016). Prokka annotation reveals the pHN7A8 plasmid (GenBank accession no. JN232517) possesses conjugative transfer machinery. Both the reference plasmid, and a representative sample of 10 IncFII pHN7A8 Inc typed plasmids carrying either ISKpn26 or ISKpn14, encoded the Tra conjugative transfer operon. The abundance of IncFII plasmids encoding ISKpn26 and ISKpn14 may be linked to efficient conjugation of IncFII host plasmids and may represent a further independent mechanism promoting the emergence of colistin resistance.

In contrast to the other IS elements investigated, IS903B was sourced from environmental sources more than the other IS elements investigated and contains the largest number of unique Inc groups. This may represent the divergence sources IS903B-containing plasmids are derived from. Concordant with these results, 37.5% (*n*=3/8) of *K. pneumoniae* samples derived from food sources from India harboured IS903B insertions in the *mgrB* gene, confirming environmental sources harbour IS elements which disrupt *mgrB* (Ghafur *et al*., 2019). Furthermore, the IS903B IS element has been detected from a bovine veterinary sample inserted into the *mgrB* gene of a *K. pneumoniae* isolate conferring colistin resistance (Kieffer *et al*., 2015). It is noteworthy, that plasmids carrying IS903B have fewer carbapenemase genes relative to the other IS elements investigated; this may reflect a clinical scenario whereby the selective pressure imposed in hospital environments engenders a situation whereby plasmids preferentially acquire carbapenemase genes in response to treatment regimens.

Whole genome sequencing (WGS) is being increasingly employed in bacterial genomics to support pathogen surveillance. The results indicate plasmid Inc groups and the country of origin for particular *mgrB* disrupting IS elements. Moreover, the relationship between IS elements and the co-carriage of carbapenemase genes and respective isolation source is exposed. Importantly however, other IS elements can also disrupt the *mgrB* gene including IS10R, ISEcp1, and ISKpn28 (Jayol *et al*., 2015; Zaman *et al*., 2018; Yang *et al*., 2020). Furthermore, IS elements can disrupt other chromosomal colistin associated genes including *crrCAB*, giving rise to colistin resistant *K. pneumoniae* (Yang *et al*., 2020). Functional studies involving IS elements in plasmid vectors transformed into colistin susceptible *K. pneumoniae* and other Enterobacteriaceae isolates will also more fully reveal the role of IS-carrying plasmids in the induction of colistin resistance. The discovery of the co-carriage of carbapenemase genes with IS elements in clinical samples indicate *K. pnuemoniae* strains are primed for resistance towards last-resort antibiotics, limiting already narrow therapeutic treatment options.

